# Understanding psychophysiological interaction and its relations to beta series correlation

**DOI:** 10.1101/322073

**Authors:** Xin Di, Zhiguo Zhang, Bharat B Biswal

## Abstract

Psychophysiological interaction (PPI) was proposed 20 years ago for study of task modulated connectivity on functional MRI (fMRI) data. A few modifications have since been made, but there remain misunderstandings on the method, as well as on its relations to a similar method named beta series correlation (BSC). Here, we explain what PPI measures and its relations to BSC. We first clarify that the interpretation of a regressor in a general linear model depends on not only itself but also on how other effects are modeled. In terms of PPI, it always reflects differences in connectivity between conditions, when the physiological variable is included as a covariate. Secondly, when there are multiple conditions, we explain how PPI models calculated from direct contrast between conditions could generate identical results as contrasting separate PPIs of each condition (a.k.a. “generalized” PPI). Thirdly, we explicit the deconvolution process that is used for PPI calculation, and how is it related to the trial-by-trial modeling for BSC, and illustrate the relations between PPI and those based upon BSC. In particular, when context sensitive changes in effective connectivity are present, they manifest as changes in correlations of observed trial-by-trial activations or functional connectivity. Therefore, BSC and PPI can detect similar connectivity differences. Lastly, we report empirical analyses using PPI and BSC on fMRI data of an event-related stop signal task to illustrate our points.

## 1. Introduction

The study of task modulated brain connectivity on functional MRI (fMRI) data is critical for understanding brain functions (Karl J Friston, 2011; Park & Friston, 2013). Unlike resting-state, where the entire scan period can be treated as a single condition (B. B. Biswal et al., 2010; B. Biswal, Yetkin, Haughton, & Hyde, 1995; Di & Biswal, 2014), for task based fMRI, multiple task conditions are typically designed within a scan run. The challenge is to estimate brain connectivity differences between different conditions. In this paper, we mainly focus on two methods that have been developed to study task modulated connectivity for task fMRI data, namely psychophysiological interaction (PPI) (K J Friston et al., 1997) and beta series correlation (BSC) (Rissman, Gazzaley, & D’Esposito, 2004).

PPI was first proposed by Friston and colleagues based on the interaction term between a physiological variable of a regional time series and a psychological variable of task design in a regression model (K J Friston et al., 1997). According to the definition of effective connectivity (Karl J. Friston, 1994), which refers to the directed effect that one brain region has on another under some model of neuronal coupling, a PPI can be regarded as a condition specific change in effective connectivity, under a simple general linear model (GLM) of interregional coupling. Thereafter, a major update was made to perform deconvolution on the time series from the seed region, so that the interaction term could be calculated at the “neuronal level” rather than at the hemodynamic response level from fMRI signals (Gitelman, Penny, Ashburner, & Friston, 2003). Later, McLaren and colleagues proposed a “generalized” PPI approach for modeling PPI effects for more than two conditions (McLaren, Ries, Xu, & Johnson, 2012). They proposed to model each task condition separately with reference to all other conditions and then compare the PPI effects between the conditions of interest, rather than to directly calculate PPI effects between the two conditions. Recently, we found that the interaction between a failure to center the psychological variable and imperfect deconvolution process may lead to spurious PPI effects (Di, Reynolds, & Biswal, 2017), and the deconvolution may be not a necessary step for PPI analysis on block-design data (Di & Biswal, 2017).

The BSC method, on the other hand, was primarily proposed for event-related designs (Rissman et al., 2004). By modeling the activations of every trial separately in a GLM, one can estimate a series of beta maps representing the activations of the series of trials. Therefore, connectivity in different task conditions can be calculated and compared in terms of correlations of trial-by-trial beta series variability. According to the distinction of functional connectivity (Karl J. Friston, 1994), which refers to the correlations in measured responses in two areas, a BSC provides a measure of functional connectivity. The relations between the BSC and PPI have not yet been clearly explained. Nevertheless, one study has suggested that BSC method is more suitable for event-related data than PPI (Cisler, Bush, & Steele, 2014). However, our recent study using a large sample did not support this conclusion (Di & Biswal, 2019).

In the current paper, we aim to provide an in-depth explanation of the PPI and BSC methods, and explain the relations and differences between these two methods. In order to do so, we need to first clarify why the PPI method always measure connectivity differences between conditions. In addition, we explain the deconvolution process implemented in the calculation of PPI, which can help to understand how PPI can measure the differences between the trial-by-trial dependency in one condition and the moment-to-moment dependency in the remaining time points. Because of this, the PPI differences between conditions and the BSC differences between conditions can, in principle, measure different aspects of task-related modulations of connectivity. We use real fMRI data of an event-related designed task to illustrate our points.

### 1.1. Modeling of task main effects

One critical point on understanding modeled effects in a model is that the interpretation of a regressor in a model depends not only on itself but also on how other effects are modeled. To explain this, we start with the modeling of the main effects of task conditions. Assume a simple task design of two conditions A and B, e.g. a flickering checkerboard condition and a fixation baseline. A regression model needs two regressors to represent the two conditions, which can be expressed in two ways. First, one can use the two regressors to represent the specific effect of each condition, i.e. using 1 to represent one condition and 0 for the other condition (Figure 1A). However, a constant term that represents the overall effect, a.k.a. intercept, is usually added in a regression model. Therefore, we only need to add one more regressor to represent the differential effect between the two conditions (Figure 1B). The two models are mathematically equivalent, because the two regressors in model 1A can be expressed as linear combinations of the two regressors in model 1B, and vice versa. However, because of the differences model strategies, the meanings of the same regressor from the two models (the first regressor from model 1A and the second regressor from model 1B) have changed. In model 1A, the regressors represent the condition specific effects. In model 1B, however, the second regressor actually represents the differential effect of conditions A and B. This is important regarding the interpretation of the estimated effects of these regressors. Mathematically, model 1B can be expressed as:

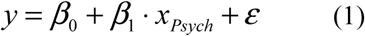

where *x_Psych_* represents the differential effects between conditions A and B, i.e. the psychological variable in the context of a fMRI data analysis. *y* represents the brain signal in a brain region or voxel. *β_0_* and *β_1_* are parameter estimates that represent the mean effect and differential effect of the two conditions, respectively.

**Figure 1.**
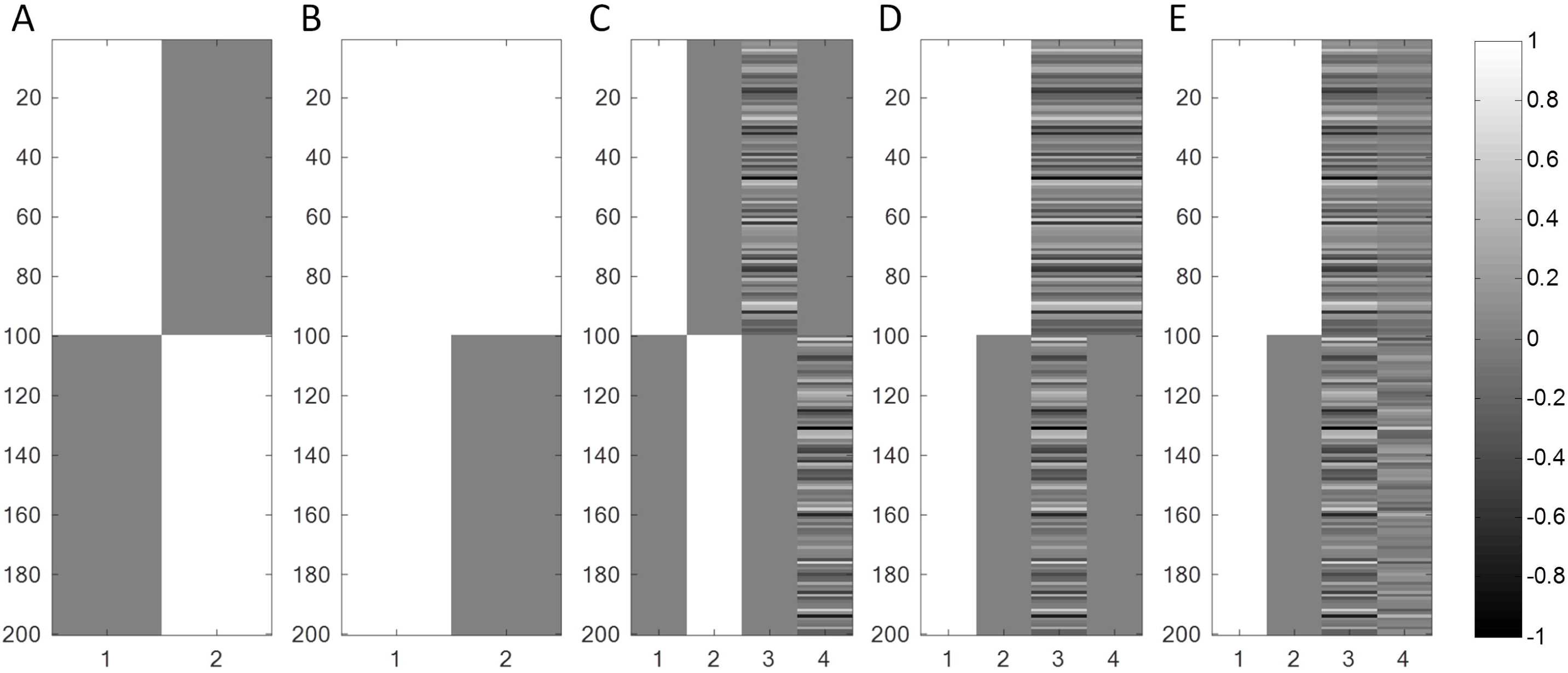
Main effects and interaction models for two experimental conditions. The main effects of two conditions can be modeled as two separate regressors (A), or modeled as the differential and mean effects of the two conditions (B). When modeling the interaction terms of the experimental condition with a continuous variable, the same two strategies could be used as C and D. E illustrates how the interaction term was changed (from D) when centering the psychological variable before calculating the interaction term. Because of the different modeling strategies, the interpretations of the regressors changed.

Another important point from equation 1 is that although *x_Psych_* is usually coded as 1 and 0 for the two conditions, the constant component in *x_Psych_* is explained by the constant term in equation 1 (see supplementary materials). Thus, whether centering the *x_Psych_* variable will not affect the effect estimate of *β_1_*, neither the interpretation of *β_1_*. *β_1_* always represents the differential effect of the two conditions.

### 1.2. Functional connectivity and connectivity-task interactions

The term functional connectivity was first defined by Friston (Karl J. Friston, 1994) as temporal correlations between spatially remote brain regions. Assuming that the functional connectivity is the same during the period of scan, e.g. in resting-state, it is straightforward to calculate correlation coefficients between two brain regions to represent functional connectivity. In a more general regression form, the model can be expressed as:

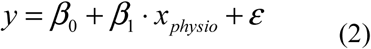

where *x_physio_* represents the time series of a seed region. *β_1_* in this case represents the regression slope, when regressing the activity of the tested voxel on a seed voxel response, i.e., a simple form of effective connectivity.

In most of task fMRI experiments, researchers design different task conditions within a scan run, so that the effect of interest becomes the differences of temporal dependencies between the conditions. We can combine equations 1 and 2 to include both the time series of a seed region (the physiological variable) and the psychological variable into a regression model. Most importantly, the interaction term of the psychological and physiological variables can also be included. For the simplest scenario of only one psychological variable (two conditions), the psychophysiological interaction model can be expressed as:

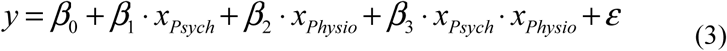

Equation 3 can be illustrated figuratively in Figure 1D. Combine the two terms with *x_Physio_*, equation 3 can be expressed as:

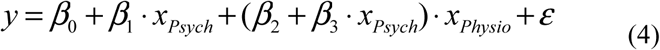

So that the relationship between the seed region *x_Physio_* and test region *y* is: *β*_2_+*β*_3_· *x_Psych_*, which is a linear function of *x_Psych_*. Therefore, a significant *β_3_* represent significant task modulations on effective connectivity.

Similar to the interpretation of task main effects, the interpretation of the PPI effect depends not only on the PPI itself, but also the on other regressors in the model. In equation 3 the main effect of time series *x_Physio_* is included. We can think about the time series main effect *x_Physio_* and interaction effect *x_Physio_ x_Psych_* as the second order counterparts of the constant effect and main effect of *x_Psych_* in equation 1. Here, adding this time series main effect affects the interpretation of the interaction term. Because the overall relationship with the seed time series has been explained by the main effect, the interaction term measures the differences of the relationships between the two conditions. We note that if *x_Physio_* main effect was not added, the interaction term could actually be calculated with each condition separately (Figure 1C). Then the third and fourth columns in Figure 1C can represent condition specific connectivity effects. Again, the same interaction terms from the two models (regressor 3 in model 1C and regressor 4 in model 1D) represent different effects.

In addition, because the main effects of *x_Physio_* and *x_Psych_* are both added in the interaction model (equation 3), the interpretation of the interaction term should refer to the demeaned version of the two variables. Because the *x_Psych_* is usually coded as 0 and 1 for the two conditions, the demeaned version of *x_Psych_* will be −0.5 and 0.5 instead. This will make the interaction term look very different (column 4 in Figure 1E compared with that in Figure 1D). However, the estimated interaction effect will be identical, because the difference between the two interaction terms is the physiological main effect, which has been taken into account in the model (For real fMRI data, however, the centering matters because the main physiological main effect interacts with the deconvolution process to produce spurious PPI effects. See Di, Reynolds, & Biswal (2017) for more details).

To better illustrate the meaning of PPI effect, we plot the PPI effect against the original time series *x_Physio_* (Figure 2). PPI can be represented as a projection of the seed time series, so that the PPI represents different relationships with the seed region in different task conditions. When the psychological variable is coded as 1 and 0 for the two conditions, the PPI represents a perfect relationship with the seed time series in the “1” condition and a smaller effect in the “0” condition, which is reflected as a horizontal line in Figure 2D. When the mean of the psychological variable is removed before calculating the interaction term, the projection rotates clockwise compared with the non-centered version (Figure 2G). However, what is reflected in the two projections are the same, which is the difference between the two conditions. In real cases, there may be positive connectivity in condition A and no connectivity in condition B, or there may be no connectivity in condition A but negative connectivity in condition B. In both cases, PPI can capture the differential connectivity effects between A and B.

**Figure 2.**
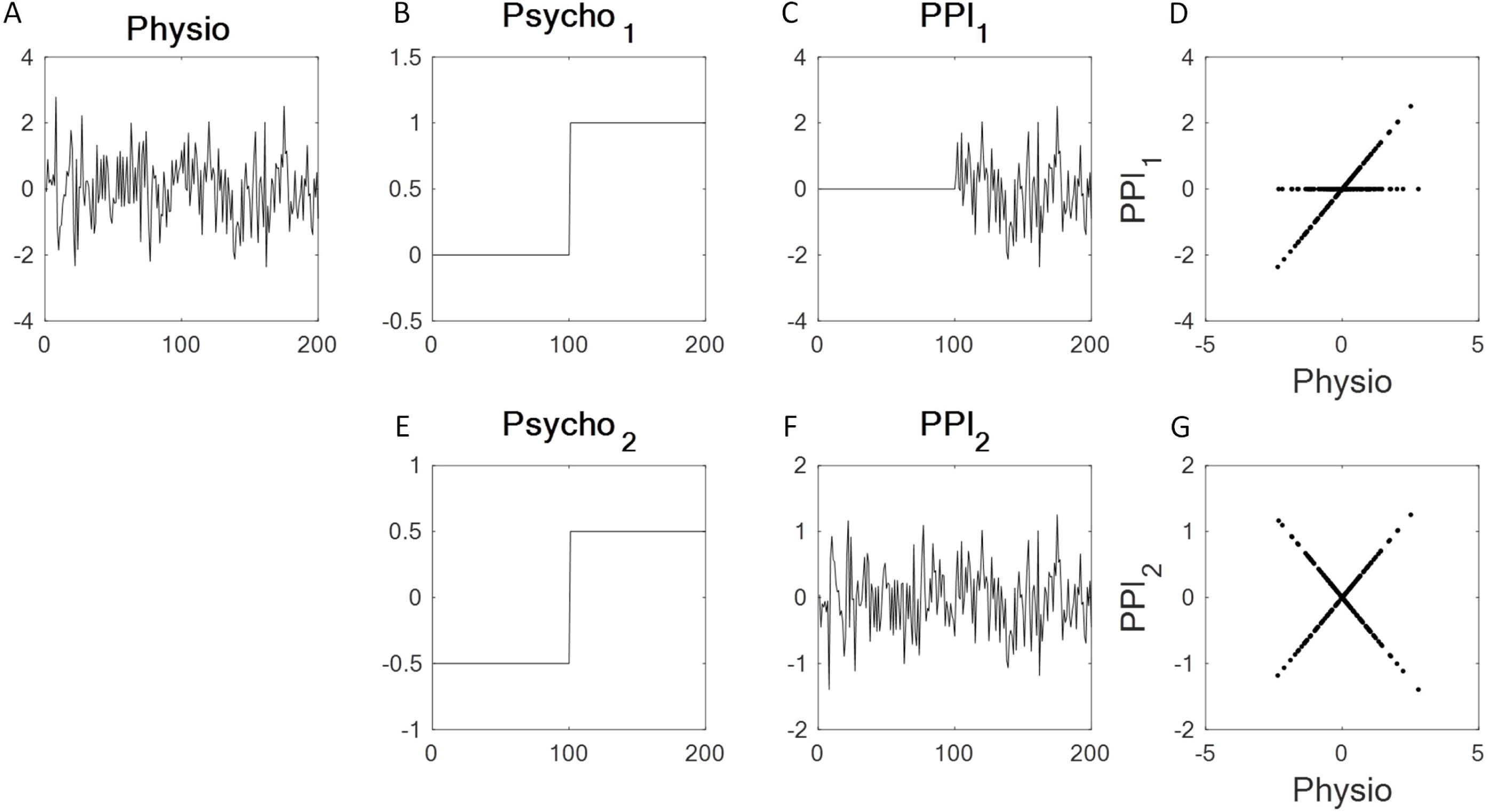
The interaction term as a projection of the continuous (physiological) variable. A continuous variable (A) is multiplied with a psychological variable (B or E) to form an interaction term (C or F), which can be plotted against the continuous variable itself (D or G). When the psychological variable is coded as 0 and 1 (B), the projection will result in a horizontal line (*y =* 0) during the 0 period and a *y = x* line during the 1 period. But usually the psychological variable is centered (E). Therefore, the projection represents *y =* - 0.5 · *x* and *y =* 0.5 · *x* lines during the two conditions, respectively.

### 1.3. More than two conditions

If we now consider multiple conditions or experimental factors, there is an opportunity to evaluate the differences in the PPI that is induced by one psychological factor, relative to another. This becomes a difference in PPIs that, mathematically, can be thought of as the interaction between the two psychological factors in mediating the interaction with the physiological fluctuations. Technically, this is a three-way interaction sometimes referred to as a psycho-psychophysiological interaction (PPPI) (Rowe, Stephan, Friston, Frackowiak, & Passingham, 2005). For example, if two sorts of face stimuli were presented (e.g., angry versus neutral) a transient increase in coupling from the amygdala to the fusiform area might be greater under the angry faces (relative to no faces), compared to the neutral faces (relative to no faces). In other words, the condition specific change in effective connectivity itself depends upon whether the stimuli were angry or neutral.

When considering modeling *n* number of conditions, the same *n* regressors are needed. Because there is always a constant term in the regression model, we actually need *n – 1* additional regressors. This is convenient for most task fMRI studies, because there is usually an implicit baseline condition, e.g. a resting or fixation condition. For event-related design, it is even difficult to explicitly define the baseline condition. Therefore, we can include all the experimental conditions, and leave the baseline condition out of the model. Because of the inclusion of the constant term, we should always keep in mind that the regressors included in the model represent the differences of between the modeled condition with respect to all the other conditions, rather than the specific effect of a condition.

Here assume a task design with conditions A and B together with a baseline condition R. The effect of interest is the differences between conditions A and B. A natural way to model the three conditions is to use two regressors to represent A and B, separately (Figure 3B and 3D). We could then calculate the interaction terms of the two psychological regressors with the seed time series, respectively. The two interaction terms represent the connectivity differences effects of A - (B + R) and B - (A + R), respectively. A contrast between these two, i.e. [A – (B + R)] – [B – (A + R)] = 2 × (A - B), can then be used to examine the differences between A and B. This strategy is usually referred to as “generalized” PPI (McLaren et al., 2012). Alternatively, one can directly contrast A with B to define a new psychological variable. It can be achieved in SPM by defining contrast value 1 to condition A, and −1 to condition B (Figure 3F). However, don’t forget the third condition R, which will be implicitly left as 0. Simply doing this is problematic, because it assumes that the relationship in the R condition is somehow between what are in the A and B conditions (Figure 3G). Because there are three conditions in total, we have to use two variables to model the differences among the three conditions. In this case, we could include one more psychological variable to represent the differential effect between the mean effect of A and B and the effect of R (Figure 3H). The interaction term of this psychological variable with the seed time series can effectively remove the differential effects of relationships between conditions A/B and condition R (Figure 3I). Therefore, if we include the PPI terms of the psychological variables 3F and 3H in the model, the effect corresponding to 3F will be equivalent to the differential effects of the PPIs corresponding to 3B and 3D. In the original paper of McLaren, it has been shown that the “generalized” PPI approach performed better than the contrast PPI. It is probably because of the neglect of the R condition. If the psychological variables are modeled correctly, the two methods should provide the same results.

**Figure 3.**
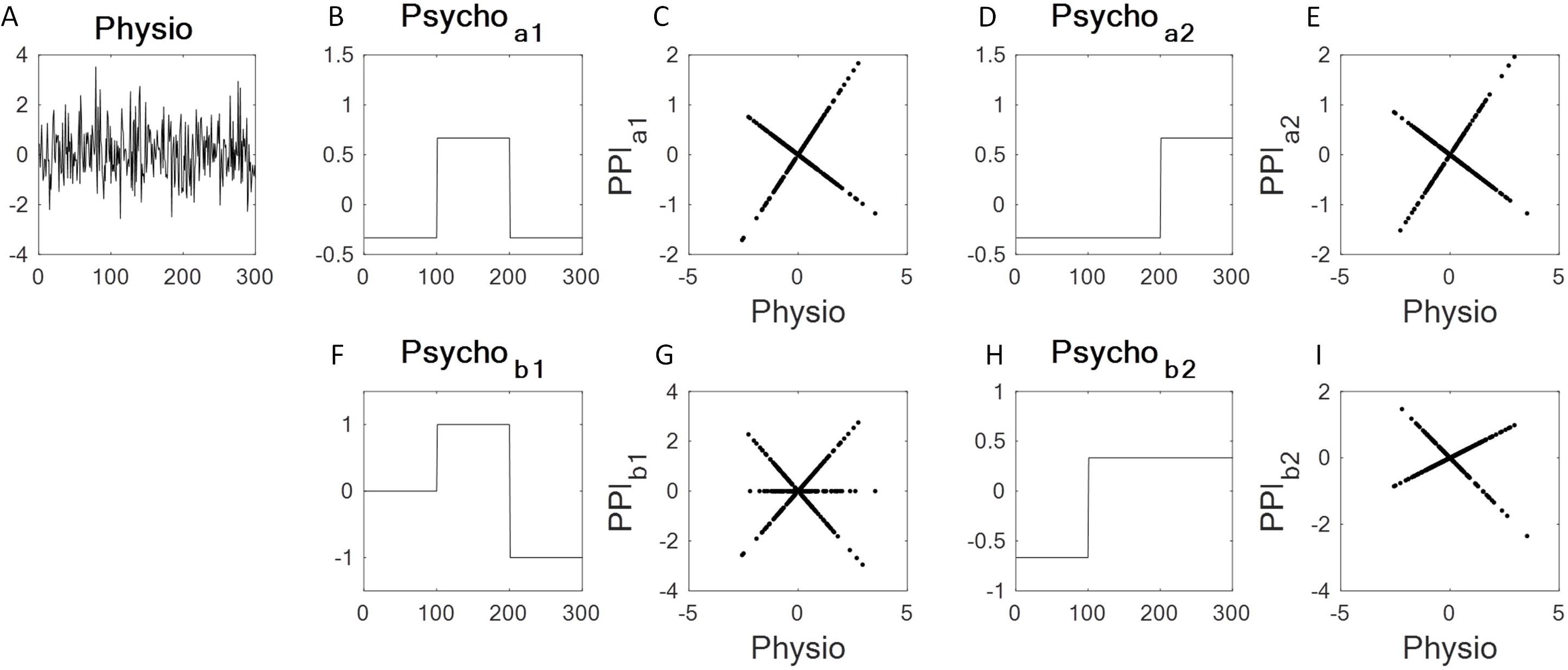
Illustrations of “generalized” PPI and contrast PPI for three conditions. Because of the inclusion of the constant term, two psychological variables are needed to model the differences among the three conditions. For the “generalized” PPI approach, the two psychological variables are demonstrated as B and D, which represent one specific condition against the other two conditions. The corresponding PPI terms were plotted against the physiological variable (A) in C and E. For the contrast PPI approach, the two psychological variables are demonstrated as F and H, which represent the differential and mean effects of the last two conditions. The corresponding PPI terms were plotted against the physiological variable (A) in G and I.

### 1.4. Block design and event-related design

So far we have divided the observations of different task conditions into different groups regardless of the orders of the observations. For fMRI, the task conditions need to be designed carefully to accommodate the properties of hemodynamic responses following neural activity changes due to task designs. There are usually two types of designs, block and event-related designs. For the block design, a task condition is broken into separate short blocks, and the blocks are repeated for several times within a scan run. In block designs, the psychological factor is usually thought of as a context (e.g., attentional set). In this setting, the PPI can be interpreted in terms of the difference in the regression slopes when regressing activity in one area on the other, under the two experimental contexts. If one interprets the regression slope has a simple measure of effective connectivity, then the PPI corresponds to the change in effective connectivity, given the experimental context or psychological factor.

For event-related design, each trial is a unit to evoke hemodynamic responses. The temporal distance between trials should be designed carefully, so that the hemodynamic response for each trial could be effectively separated (Dale, 1999; K J Friston et al., 1998). The psychological variable for event-related design is modeled as a series of impulse function at the onset of the trials with remaining time points as 0. Mathematically, exactly the same interpretation holds for event related designs; however, here, the event is very short lived and cannot be interpreted as a context. A PPI in event related designs reflects a transient increase (or decrease) in the effective connectivity (in a trial-by-trial manner) between two brain areas that can be attributed to the stimulus for action associated with the event in question. For example, a visual stimulus may transiently enhance the coupling between the amygdala and fusiform area, in relation to baseline coupling in the absence of a stimulus.

### 1.5. Convolution and deconvolution

One important aspect of fMRI is the asynchrony between the (hypothetical) neuronal activity and the observed blood-oxygen-level dependent (BOLD) signals. A trial or short event elicits transient neural activity that is typically treated as an impulse function, and further gives rise to a delayed hemodynamic response, which is usually called hemodynamic response function (HRF) (Figure 4A). If there are designed trials or hypothetical neural activity, the expected BOLD responses can be calculated as a convolution of the design or neural activity time series with the HRF. Because the fMRI data are discrete signals, the convolution can be converted into a multiplication of the neuronal signal with a convolution matrix corresponding to the HRF. Using *z* to represent variables at the neuronal level, and *x* to represent variables at the BOLD level, the convolution can be expressed as:

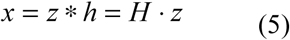

where * represents the convolution process, · represents matrix multiplication, *h* is the HRF, and *H* represents the matrix form of *h*. Each column of *H* represents a HRF with a different start point (Figure 4B). Therefore, the multiplication of a neural time series *H* with *z* can be understood as a summation of the hemodynamic responses of *z* at every time point.

**Figure 4.**
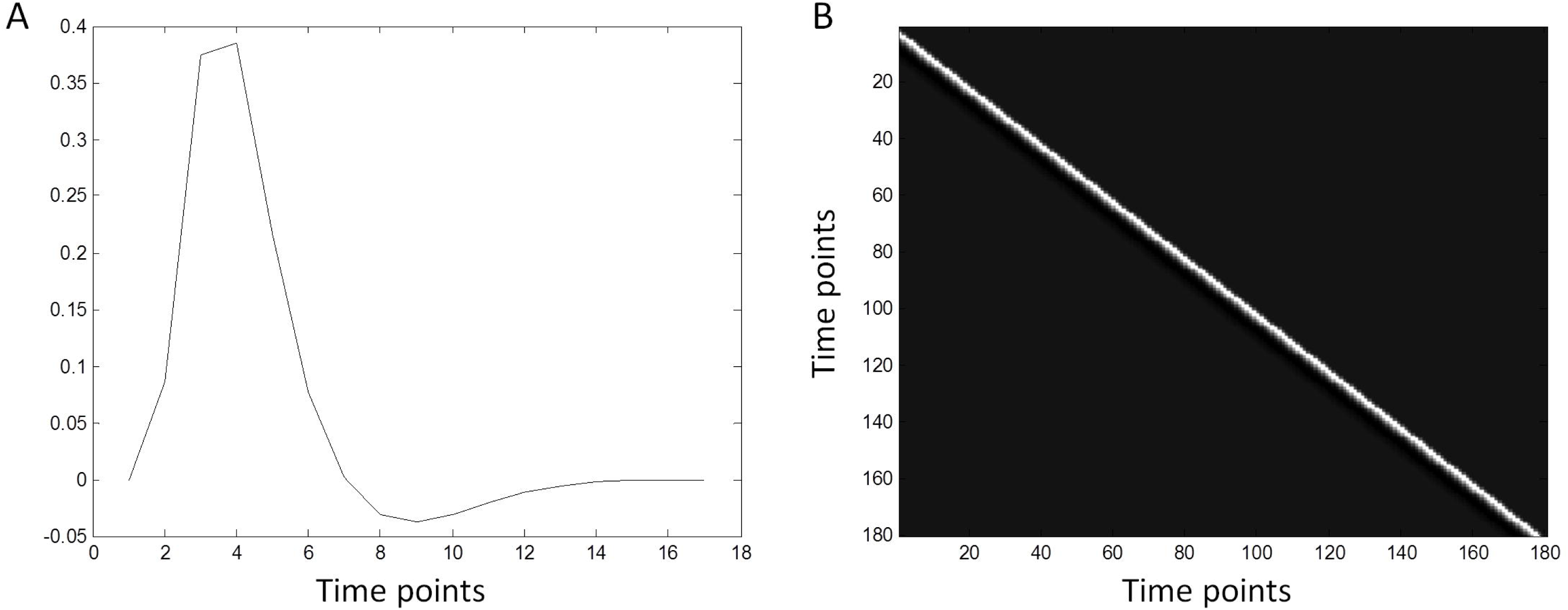
Hemodynamic response function *h* (A) and its corresponding convolution matrix *H* (B).

For fMRI, we typically hypothesize that an experimental manipulation will evoke immediate neural response (relative to the time scale of BOLD responses). The expected BOLD responses to the experimental manipulations could then be represented as the convolution of the psychological variable *z*_Psych_ (a box-car function or a series of impulse functions) with the HRF. Thus, the BOLD level prediction variable *x_Psych_* can be calculated from *z_Psych_* as the following:

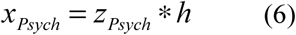

On the other hand, through fMRI we have a time series of a region *x_Physio_*, which is already at the BOLD level. Therefore, we can directly calculate the interaction term by multiplying *x_Physio_* with *x_Psych_*.

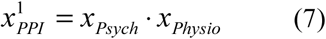

This is how PPI was calculated when the method was originally proposed (K J Friston et al., 1997). The limitation of this approach is that it calculates the interaction at the BOLD level, but the real interaction would happen at the “neuronal” level.

Given the BOLD level time series *x*, we can perform the inverse of convolution, i.e. deconvolution, to recover the time series *z* at the neuronal level from equation 5. However, the *H* matrix is a square matrix, so that deconvolution cannot be simply solved by inverting the *H* matrix. In addition, for practical deconvolution problem like the fMRI signals, there are always noises in the recorded signals that need to be taken into account. Therefore, the deconvolution problem has to solve the following model with a noise component ε

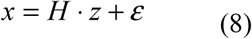

Because *H* cannot be directly inverted, some computational methods like regularization are needed to reliably obtain *z*. In SPM, it additionally substitutes *z* with Discrete Cosine Series, so that the estimation of temporal time series was transformed into frequency domain (Gitelman et al., 2003). And the regularization is applied to specific frequency components.

A seed time series *x_Physio_* can be deconvolved to the neuronal level time series *z_Physio_* and multiplied with the neuronal level psychological variable. The interaction term can then be convolved back into BOLD level:

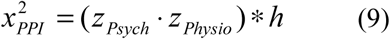

*x*^1^*_PPI_* and *x*^2^*_PPI_* are not mathematically equivalent. The later one is more appropriate to describe neural interactions. However, empirically, the PPI terms calculated with the two ways could be very similar for block designs (Di & Biswal, 2017). In addition, deconvolution is an ill-posed problem, and relies on sophisticated computational techniques, which may not work well in some circumstances. Therefore, it has been suggested that at least for block design, deconvolution may not be necessary (Di & Biswal, 2017; O’Reilly, Woolrich, Behrens, Smith, & Johansen-Berg, 2012). The deconvolution approach may still be important and necessary for event-related design.

### 1.6. Beta series correlations

BSC is based on a simple idea of calculating correlations of trial-by-trial variability of brain activations. Instead of modeling different task conditions, BSC models every trial’s activation to obtain a beta map for each trial. For each trial, an impulse function at the trial onset is defined and convolved with HRF. Therefore, a GLM for BSC analysis has the same number of regressors as the number of trials plus a constant term. The model can be expressed as the following:

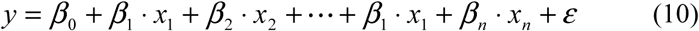

where *n* represents the number of trials, and *x_n_* represents the modeled response of the trial *n*. The model can be expressed in a matrix form:

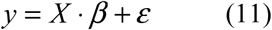

where *β* represents a vector of *βs* corresponding to the activations of different trials (plus a *β*_0_ for the constant term). Figure 5 shows examples of the design matrix *X*. One can then calculate cross-trial correlations of the beta values between regions to represent functional connectivity. Since there are usually more than one experimental condition, the beta series can be retrospectively grouped into different conditions, and the beta series correlations can be compared between the conditions.

**Figure 5.**
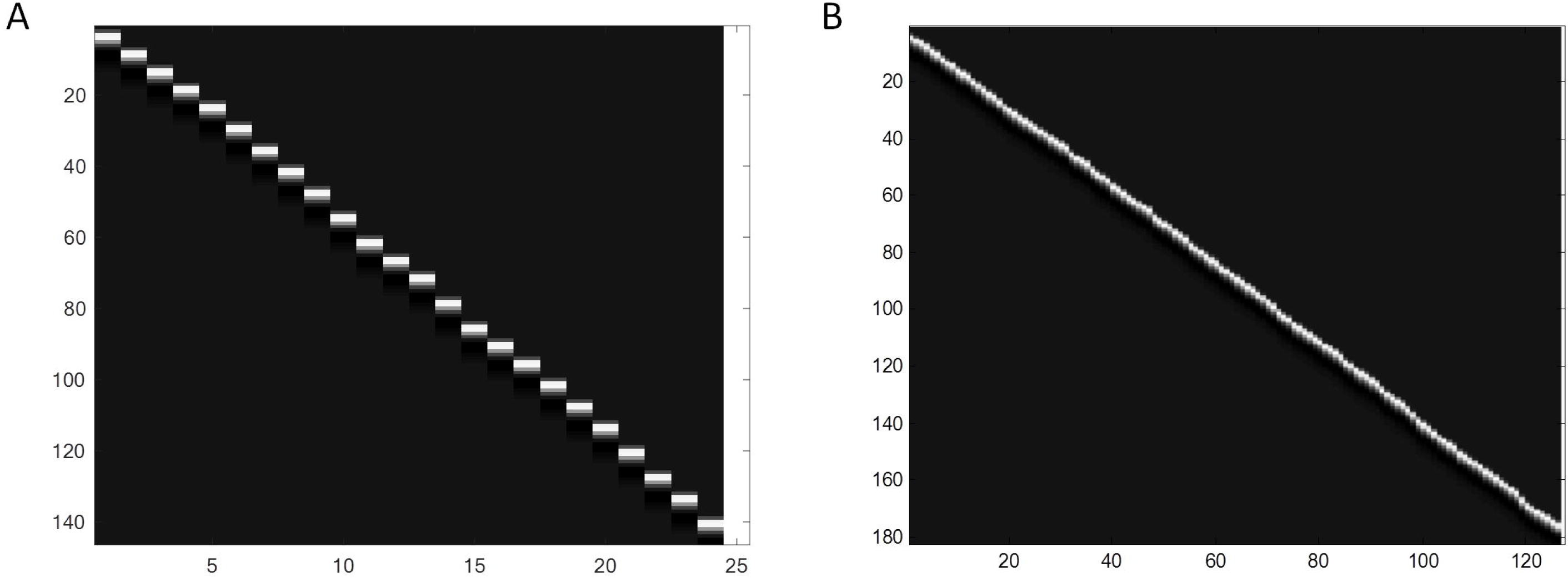
Example design matrices for beta series correlation (BSC) analysis for a slow event-related design (Flanker task) (A) and a fast event-related design (Stop signal task) (B). Each regressor (column) except the last one represents the activation of a trial, while the last column represents the constant term. The sampling time is 2 s for both of the designs. The intertrial intervals for both the designs were randomized to optimize the estimations of hemodynamic responses. The mean intertrial intervals are 12 s for the Flanker task and 2.5 s for the Stop signal task, which result in 24 trials and 126 trials, respectively.

The hemodynamic response typically reaches the peak at 6 s after trial onset and returns back to the baseline after about 15 s. To avoid overlaps of hemodynamic responses between trials, some event-related experiments use slow designs with intertrial interval usually greater than 10 s. Figure 5A demonstrates a beta series GLM for a slow event related design from a Flanker task (Kelly, Uddin, Biswal, Castellanos, & Milham, 2008). Considering a typical sampling time of 2 s for fMRI, the design matrix of Figure 5A can be reliably inverted (24 trial regressors vs. 146 time points). However, fast event-related design is becoming more standard, because of its efficiency of maximizing experimental contrasts (Dale, 1999; K J Friston et al., 1998). The intertrial interval could be close to the sampling time of fMRI for some designs. Figure 5B demonstrated a beta series GLM for a fast event-related design from a stop signal task (Di & Biswal, 2019). In this case the mean intertrial interval is 2.5 s. It can be seen that the number of regressors becomes closer to the number of time points (126 trial regressors vs. 182 time points). This matrix cannot be reliably inverted using ordinary least squares (OLS) method, and some sophisticated computational methods may be helpful to resolve the problem, e.g. using regularization or modeling a single trial against all other trials to reduce the number of regressors (Di & Biswal, 2019; Mumford, Turner, Ashby, & Poldrack, 2012).

The beta values in the beta series model typically represent BOLD level activations at each trial. However, in a case when the trials are presented at every time point, the beta series model becomes exactly the same as the convolution matrix in Figure 4B. The beta series model estimates transient neuronal responses specifically at each trial onset, while the deconvolution model estimates neuronal responses at every time point. This suggests a link between the beta series model and deconvolution.

### 1.7. The relations between PPI and BSC

As described in previous sections, the BSC method estimates neuronal activity for each trial, and computes trial-by-trial correlations between brain regions. The correlations reflect functional connectivity of a task condition. The PPI, on the other hand, measures connectivity differences as coded by a psychological variable. Therefore, a beta series correlation in one condition is not comparable to a simple PPI effect. However, for task fMRI what is usually of interest are the differences between task conditions. Considering the same task design with experimental conditions A, B, and a baseline R, as has been explained, both contrast PPI and “generalized” PPI can measure effective connectivity differences between A and B. On the other hand, BSCs can be directly compared between the conditions A and B. Therefore, in theory the BSC may be sensitive to the changes in coupling tested for by PPI analyses. However, the two analyses are not measuring the same sort of thing, because a PPI is effectively a change in regression slope or parameter of a model of effective connectivity. In contrast, differences in BSCs are changes in the statistical coupling or functional connectivity. This is important because one can have a change in a correlation without a change in effective connectivity. This can occur when the random noises in one experimental condition differ from the other.

Although theoretically PPI and BSC could detect the same task modulated connectivity, the results of PPI and BSC on real fMRI data may not be identical. Several factors may contribute to the differences. The first is the different approaches to deconvolution. The deconvolution method implemented in SPM uses Discrete Cosine Series to convert the temporal domain signal into frequency domain, and then applies regularization on the frequency domain to suppress high frequency components in the signals. For BSC method, if it is a slow event-related design, single-trial activations can be directly estimated from the GLM. For a fast event-related design, some regularization methods or complicated modeling strategies are required to estimate the beta series (Mumford et al., 2012). The efficiency and reliability of these mentioned methods are difficult to determine and compare. And it may depend on the intertrial intervals of a design (Abdulrahman & Henson, 2016; Mumford, Davis, & Poldrack, 2014; Visser et al., 2016), or different brain regions due to different amount of HRF variability (Handwerker, Ollinger, & D’Esposito, 2004).

Another difference may be the different statistical measures of connectivity. By using a regression model PPI essentially measures the differences in slope between conditions. On the other hand, BSC typically uses correlation coefficients. It is still largely unknown how the variability of BOLD signals changes in different task conditions (Duff, Makin, Cottaar, Smith, & Woolrich, 2018), which may influence PPI measures. For BSC, one can choose different measures of connectivity, e.g. Pearson’s product-moment correlation, Spearman’s rank correlation, covariance, or even similar beta series-by-task interaction as PPI under a regression model to estimate connectivity differences. However, it is still an open question about which method is optimal for the purpose of connectivity estimations.

### 1.8. An empirical demonstration

In summary, PPI analyses allow one to test for condition specific differences in (linear) effective connectivity between two areas. Furthermore, by comparing PPIs associated with different conditions one can test for high order context sensitive changes in coupling. An obvious but interesting hypothesis here is that the presence of such high order interactions might manifest as changes in functional connectivity between two experimental conditions. One can access these changes in functional connectivity by leveraging the trial-by-trial variability in evoked responses with a BSC. One can then simply look at the differences in these correlations (i.e., functional connectivity) to see if they identify the same pairwise connections detected by “generalized” PPI analysis. We will demonstrate this using empirical data in an event related setting.

We analyzed an fMRI dataset with a fast event-related designed stop signal task. There were two experimental conditions (Go trials and Stop trials) in addition to an implicit baseline. The connectivity differences between the Stop and Go conditions have been reported previously (Di & Biswal, 2019). Here, we report connectivity measures with PPI and BSC for simple conditions and condition differences. The PPI and BSC results are also compared with resting-state functional connectivity to better illustrate their relations to each other. Secondly, we will show that the “generalized” PPI approach and direct contrast PPI approach provide identical measures of effective connectivity differences between conditions. Lastly, we will compare different correlation measures for BSC analysis, i.e. Pearson’s correlation, Spearman’s correlation, covariance, and beta series-by-task interaction.

## 2. Materials and methods

### 2.1. Dataset and designs

In our previous paper, we have reported the PPI and BSC results of connectivity differences between the Stop and Go conditions (Di & Biswal, 2019). In the current paper, we used the same data to illustrate how different PPI models could give rise to the same results and how the PPI and BSC methods can be similar or different. This dataset was obtained from the OpenfMRI database (accession #: ds000030). Only healthy subjects’ data were included in the current analysis. After removing subjects due to large head motion, a total of 114 subjects were included in the current analysis (52 females). The mean age of the subjects was 31.1 years (range from 21 to 50 years). In the stop signal task, the subjects have to indicate the direction (left or right) of an arrow presented in the center of the screen. For one fourth of the trials, a 500 Hz tone was played shortly after the arrow, which signaled the subjects to withhold their response. In a single fMRI run, there were 128 trials in total in total, with 96 Go trials and 32 Stop trials. The task used a fast event-related design, with a mean intertrial interval of 2.5 s (range from 2 s to 5.5 s). For a subset of 103 subjects, we also analyzed their resting-state fMRI data. The exclusion of additional subjects were due to large head motions in either the resting-state run or other task runs that were not included in this paper.

The fMRI data were collected using a T2*-weighted echoplanar imaging (EPI) sequence with the following parameters: TR = 2000 ms, TE = 30 ms, FA = 90 deg, matrix 64 × 64, FOV = 192 mm; slice thickness = 4 mm, slice number = 34. 184 fMRI images were acquired for each subject for the stop signal task, and 152 images were acquired for the resting-state run. The T1 weighted structural images were collected using the following parameters: TR = 1900 ms, TE = 2.26 ms, FOV = 250 mm, matrix = 256 × 256, sagittal plane, slice thickness = 1 mm, slice number = 176. More information about the data can be found in (Poldrack et al., 2016).

### 2.2. FMRI preprocessing

The fMRI image processing and analysis were performed using SPM12 (v6685) (http://www.fil.ion.ucl.ac.uk/spm/) and scripts in MATLAB R2013b environment (https://www.mathworks.com/). The anatomical image for each subject was first segmented, and normalized to standard MNI (Montreal Neurological Institute) space. The first two functional images were discarded, and the remaining images were realigned to the first image, and coregistered to the subject’s own anatomical image. The functional images were then transformed into MNI space by using the deformation images derived from the segmentation step, and were spatially smoothed using a 8 mm FWHM (full width at half maximum) Gaussian kernel.

### 2.3. PPI analysis

The first step of PPI analysis is to build a voxel-wise GLM for task activations. In the current analysis, the Go and Stop conditions were modeled separately as series of events (trials). In SPM, the durations of events are usually set as 0 to reflect their impulsive nature. But for PPI analysis, the time series are up-sampled (16 times by default) after deconvolution. If the duration is set as 0, then the neuronal level psychological variable only has a time bin of one with duration of TR/16, leaving all other time bins as 0. This may be problematic when multiplying this psychological variable with the deconvolved seed time series. Considering that the calculated PPI term will be convolved back with HRF, which resembles a low pass filtering, the effects of trial duration may not be that significant. In the previous analysis, we set the duration to 1.5 s, which represented the actual duration of the trial. We have also shown in the supplementary materials that setting the event duration as 0 produce very similar results as those with 1.5 s duration. In the results section, we report results with 1.5 s duration. In addition to the two event-related task regressors, 24 head motion regressors and one constant regressor were also included in the GLM model. After model estimation, the times series from 164 ROIs were extracted. The head motion, constant, and low frequency drift effects were adjusted during the time series extraction. These 164 ROIs were adopted from previous studies (Di & Biswal, 2019) to represent whole brain coverage. The following connectivity analyses of PPI and BSC were performed on ROI-to-ROI basis.

The PPI terms were calculated using the two different approaches. For the “generalized” PPI approach, we used the contrasts [1 0] and [0 1] to define the two psychological variables representing the Go and Stop conditions, separately. The PPI terms were then calculated using the deconvolution method. The calculated PPI terms were combined together with the original model to form a new GLM for PPI analysis:

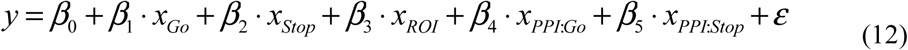

This model included one constant term, two regressors of task activations, one regressor of the ROI time series, and two regressors of PPIs. Here the differential effect of the two PPI terms, *β_5_* – *β_4_*, represents the higher order task modulated connectivity (i.e. PPPI) between the Stop and Go conditions. Because the dependent variable *y* is also a ROI time series, where the head motion effects have already been removed, the head motion regressors were no longer included in the PPI models.

For the second contrast PPI approach, we defined the differential and mean effects of the Stop and Go conditions using the contrasts [-1 1] and [1/2 1/2], respectively. After PPI calculation using the deconvolution method, the GLM for the contrast PPI analysis was defined as follow:

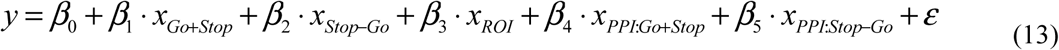

The estimates of *β_5_* were used for group level analysis to present connectivity differences between the Stop and Go conditions.

For each subject, the PPI models were built for each ROI, and were fitted to all other ROIs. The beta estimates or contrasts of interest were calculated between each pair of ROI, which yielded a 164 by 164 matrix for each effect. The matrices were transposed and averaged with the original matrices, which yielded symmetrical matrices. This procedure ensures that the PPI matrices can be compared to the (symmetrical) undirected functional connectivity matrices furnished by BSC. One sample t test was performed on each element of the matrix for an effect of interest. False discovery rate (FDR) correction was used at p < 0.05 to identify statistical significant effects in a total of 13,366 effects (*164 x (164 – 1) / 2*).

### 2.4. Beta series analysis

As has been shown in our previous paper (Di & Biswal, 2019), modeling all trials together in a single model could not work for the beta series analysis. Therefore, we only reported the results from the single-trial-versus-other-trials method (Mumford et al., 2012). We first built a GLM for each trial, where the first regressor represented the activation of the specific trial and the second regressor represented the activations of all the remaining trials. The 24 head motion parameters were also included in the GLMs as covariates. The duration of events was set as 0. After model estimation, beta values of each ROI were extracted for each trial. The beta series of each ROI were sorted into the two conditions, and the functional connectivity measures across the 164 ROIs were calculated. In our previous work, we used Spearman’s rank coefficients to avoid Gaussian distribution assumption of the beta series or spurious correlations due to outliers. In the current analysis, we also calculated Pearson’s correlation coefficients, covariance, and beta series-by-task interaction to examine whether these two measures may give more reliable estimates of connectivity. The whole beta series (Go and Stop together) of a ROI were first z transformed, and then the correlation coefficients and covariance matrices were calculated for each subject. The Pearson’s and Spearman’s correlation matrices were transformed into Fisher’s z matrices. Similar to PPI, we calculated beta series-by-task interaction for each ROI, and built a GLM including the raw beta series main effect, task main effect, their interaction, and a constant term. The GLM was used to *βs* corresponding to the interaction term were used to represent task modulated connectivity. The β matrices were transposed and averaged with the original matrices, resulting in symmetrical matrices.

For a single condition, the mean of Fisher’s z values or covariance values were averaged across subjects. Paired t tests were also performed to compare the differences between the two conditions at every element of the matrix. For the beta series-by-task interaction analysis, one sample t test was used in the group level analysis. A FDR correction at p < 0.05 was used for all the measures to identify statistical significant effects.

### 2.5. Resting-state connectivity

A voxel-wise GLM was first built for each subject, which included 24 head motion regressors and on constant term. After model estimation, the times series from the 164 ROIs were extracted, adjusting for the head motion, constant, and low frequency drift effects. For each subject, a Pearson’s correlation coefficient matrix was calculated across the 164 ROIs. The matrices were transformed into Fisher’s z matrices, and averaged across subjects.

## 3. Results

### 3.1. “Generalized” PPI vs. BSC

We first show task modulated connectivity matrices calculated from “generalized” PPI and BSC (Figure 6). To illustrate overall patterns, the matrices were not thresholded. Consistent with our explanations of the differences between PPI and BSC, the simple PPI effects of a condition look very different from those calculated from BSC. For the Go condition, the PPI matrix (shown in Figure 6A, corresponding to *β_4_* in equation 12) had increased connectivity between visual and sensorimoter regions and between cerebellar and sensorimotor regions, and decreased connectivity within visual areas compared with the other conditions. In contrast, the BSC matrix (Figure 6D) of the Go condition showed square like high correlations along the diagonal, which represent higher functional connectivity within each functional modules. The correlation between the PPI and BSC matrices of the Go condition was only 0.17 (Figure 6G). Similarly, the PPI (Figure 6B, corresponding to *β_5_* in equation 12) and BSC (Figure 6E) matrices of the Stop condition also looked different, with a small correlation of −0.20 (Figure 6H). However, despite the differences in single conditions, the differential effects between the Stop and Go conditions were similar between the PPI (Figure 6C) and BSC (Figure 6F) methods. The correlation between the two matrices was 0.73 (Figure 6I), which has been reported previously (Di & Biswal, 2019).

**Figure 6.**
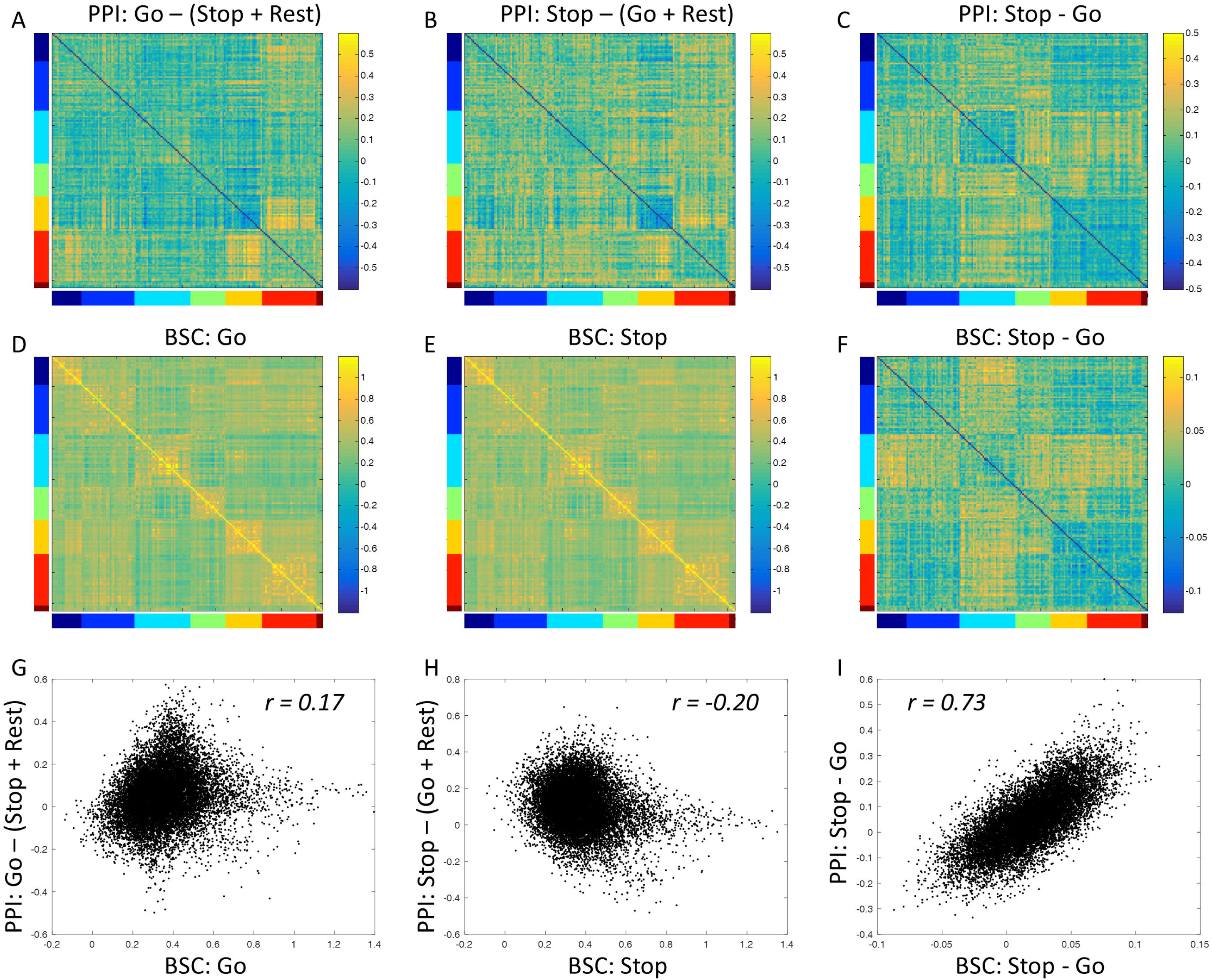
Psychophysiological interaction (PPI) and beta series correlation (BSC) results from the stop signal task. The top row showed the PPI matrices using the “generalized” PPI model, where the Go condition and Stop condition were modeled separately. The middle row showed correlation matrices using the beta series method. The bottom row shows the scatter plots between the PPI and BSC matrices of the corresponding columns. The right-side color scales of all matrices were made sure to be positive and negative symmetrical, but the range was adjusted based on the values in each matrix. The left and bottom color bars indicate the seven functional modules, including cerebellar, cingulo-opercular, default mode, fronto-parietal, occipital, sensorimotor, and emotion modules from dark blue to dark red.

The BSC matrices of both the Stop and Go conditions looked very similar to each other, and were indeed similar to resting-state correlations. To confirm this, we analyzed the resting-state fMRI data from a subset of 103 subjects, and calculated resting-state functional connectivity matrix (Figure 7A). The correlations between the resting-state connectivity and the BSC matrices of the Go and Stop conditions were 0.91 (Figure 7B) and 0.92 (Figure 7C), respectively.

**Figure 7.**
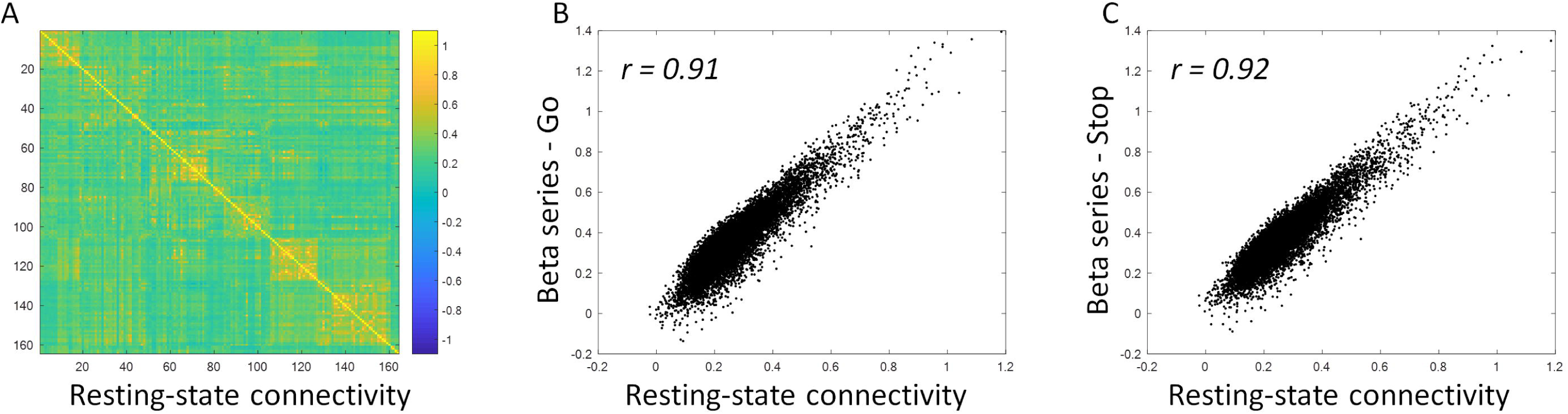
Resting-state functional connectivity matrix (mean Fisher’s z) from a subset of 103 subjects (A), and its relationship with the beta series correlations (BSC) matrices of single conditions (B and C).

### 3.2. “Generalized” PPI vs. contrast PPI

We also performed PPI analysis using the contrast PPI approach, i.e. modeling the mean and differential effects of the Go and Stop conditions, respectively. The mean PPI effects of the Go and Stop conditions (Figure 8A, corresponding to *β_4_* in equation 13) looked similar to the specific effects of each condition from the “generalized” PPI approach (Figure 6A and 6B). Most interestingly, consistent with our explanation, the differential effects of the Stop and Go conditions (Figure 8B, corresponding to *β_5_* in equation 13) turned out to be identical to the contrast of Stop and Go PPI effects from the “generalized” PPI model (Figure 6C). This can be confirmed by showing the scatter plot between the two matrices (Figure 8C), which demonstrated as a straight line. It should be noted that the effects of the “generalized” PPI (contrast values) were as two times as the effects of the contrast PPI. This has been explained in section 1.3 that the contrast between two “generalized” PPI effects A and B represent the effect of 2 × (A - B).

**Figure 8.**
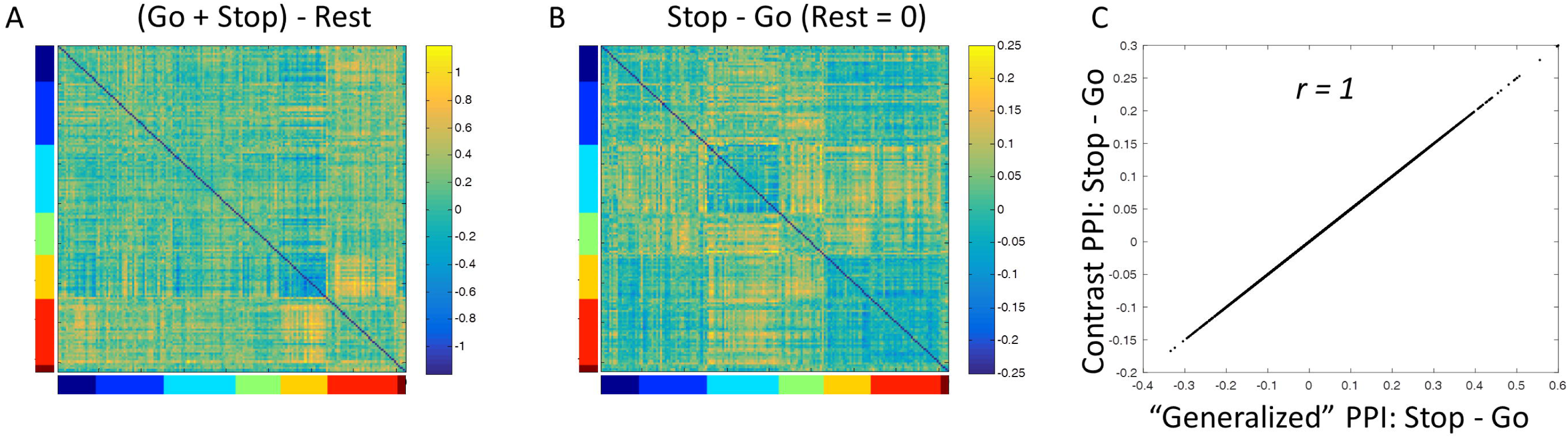
Psychophysiological interaction (PPI) results of the mean effects (A) and differential effects (B) between the Stop and Go conditions using contrast PPI. C shows the relationship between the Stop – Go contrasts calculated from the contrast and “generalize” PPI methods.

The mean effects of the Go and Stop conditions, which reflect task modulated connectivity related to general task execution, have not been reported previously. We performed one sample t test on every element of the matrix. Statistical significant effects were thresholded at p < 0.05 (FDR corrected) and visualized using BrainNet Viewer (Xia, Wang, & He, 2013) (Figure 9). It clearly shows that there was reduced effective connectivity within the visual areas, and increased connectivity mainly between visual regions and sensorimotor regions and between visual regions and other brain regions such as cingulo-opercular regions.

**Figure 9.**
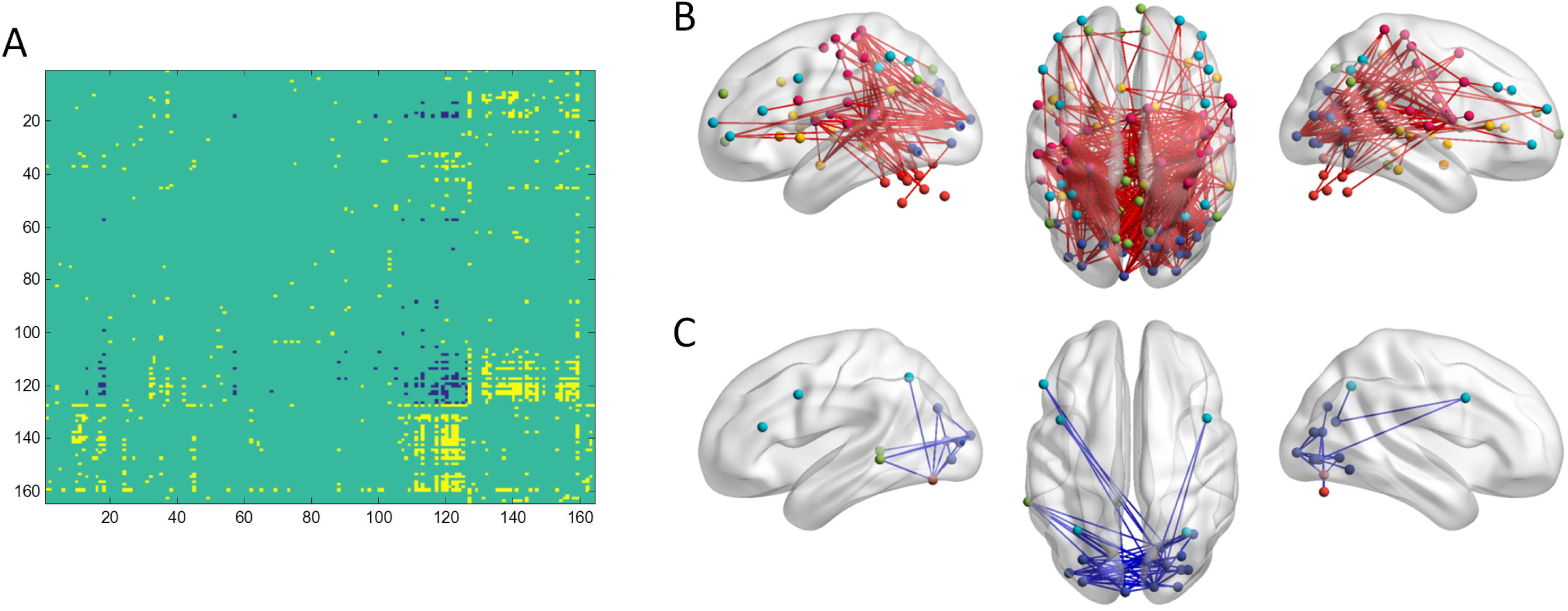
Mean PPI effects of the Go and Stop trials compared with the implicit baseline. A shows the thresholded PPI matrix at p < 0.05 of FDR (false discovery rate) correction. Yellow represents positive PPI effects, while blue represents negative effects. The color bars indicate the seven functional modules, including cerebellar, cingulo-opercular, default mode, fronto-parietal, occipital, sensorimotor, and emotion modules from dark blue to dark red. B and C show the positive and negative effects on a brain model using BrainNet Viewer.

### 3.3. Different BSC measures

Lastly, for the contrast of Stop vs. Go where the PPI and BSC methods yielded similar results, we compared different correlation measures for the BSC analysis (Figure 10). When comparing the four measures of Spearman’s correlation, Pearson’s correlation, covariance, and beta series-task interaction, Pearson’s correlation produced the largest number of significant effects, while covariance only showed one significant positive effect and one significant negative effect. However, even the results from Pearson’s correlation showed less significant results than the PPI model.

**Figure 10.**
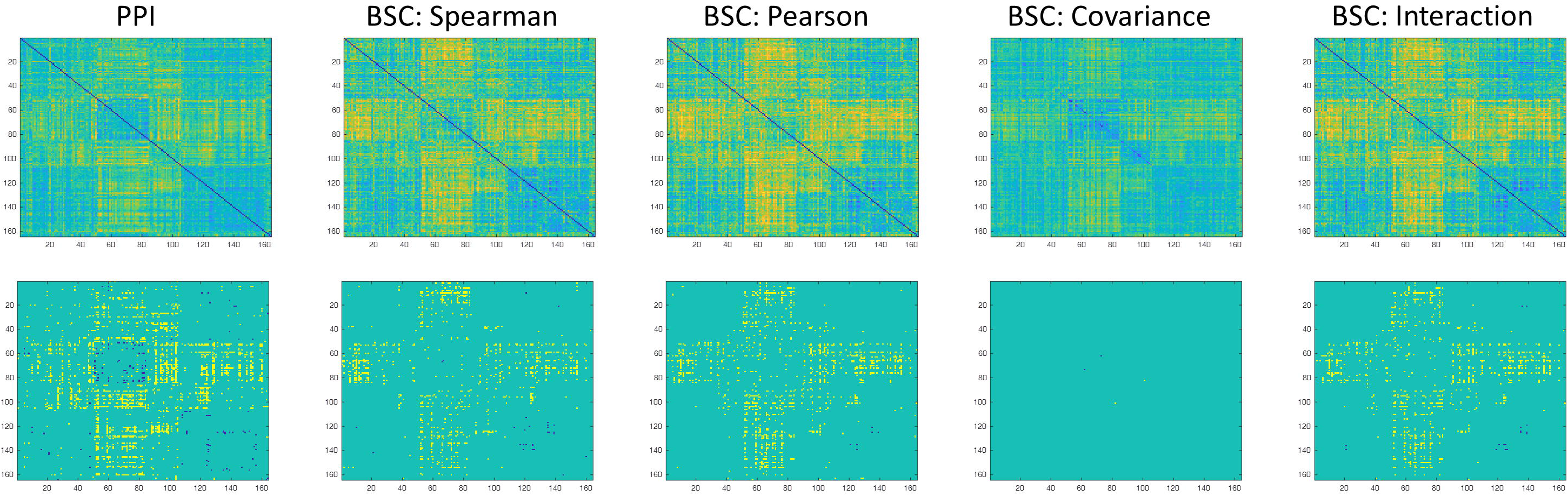
Unthresholded (upper row) and thresholded (lower row) matrices of task modulated connectivity between the Stop and Go conditions estimated by different methods. A p < 0.05 of false discovery rate (FDR) correction was used to threshold each matrix. The color scales of all matrices were made sure to be positive and negative symmetrical. But the range was adjusted based on the values in each matrix.

## 4. Discussion

In the current paper, we have explained that because the inclusion of the physiological variable in the PPI model, a PPI effect always represents the differences of regression parameters between conditions. In an event-related setting, a PPI reflects effective connectivity differences between those elicited by transient events and those during the baseline condition. In contrast, BSC measures functional correlations elicited by series of transient events in a specific task condition. However, when comparing between conditions, BSC should in principle yield similar estimates as PPI differences (i.e. PPPI effects). The results of PPI and BSC analyses on fMRI data of an event-related designed stop signal task agree with our theoretical explanation of the two methods. Firstly, the BSCs of the Go and Stop conditions were very similar to each other, and were also highly correlated with the functional connectivity in resting-state. In contrast, the PPIs of the Go and Stop condition from the “generalized” PPI approach tuned out to be very different from those from the BSC analysis. Secondly, when contrasting the Stop and Go conditions, the PPI matrices and BSC matrices were very similar, which reflected the connectivity differences between the conditions. Lastly, consistent with our mathematical explanation, direct contrast PPI showed exactly the same results as “generalized” PPI when the conditions were modeled properly.

As explained in the introduction, the correlations of trial-by-trial variability measured by BSC reflect the absolute level of functional connectivity in a condition. Interestingly, we found that the BSCs for the Go and Stop conditions were very similar to each other, and were also similar to what we typically observed in resting-state. This is consistent with the observation that the moment-to-moment correlations in many different task conditions are very similar (Cole, Bassett, Power, Braver, & Petersen, 2014). This suggests that the absolute correlations between brain regions in any task conditions are highly contributed by the correlations of spontaneous neural activity (B. Biswal et al., 1995), or other common factors that could give rise to high correlations, e.g. common anatomical connectivity (Honey et al., 2009), common neurovascular responses (Hillman, 2014; Sivakolundu et al., 2019), physiological noises (Weissenbacher et al., 2009), or head motion (Power, Barnes, Snyder, Schlaggar, & Petersen, 2012; Van Dijk, Sabuncu, & Buckner, 2012). Nevertheless, the similarity of absolute correlations in many different task conditions including resting-state makes them lack of interest in terms of understanding brain functions. Practically, it would be more informative and valuable to examine whether connectivity is modulated by a task manipulation than to only look at the absolute value of the connectivity.

It is noteworthy that in equation 3, *β_2_* represents task independent co nnectivity between the seed region *x_Physio_* and test region *y*, after covarying task dependent connectivity and task activation. Further, given the linear function of the relation between *x_Physio_* and *y* in equation 4, i.e. *β_2_* + *β_3_* · *x_Psych_*, one can also recover functional connectivity between the two in a specific task condition encoded in *x_Psych_*. But one should be cautious that due to centering and deconvolution process, the values in *x_Psych_* to represent one task condition will no longer be 1 (usually smaller than 1). Simply using *β_2_* + *β_3_* will over emphasize task modulated connectivity, and cannot be a good estimate of task specific connectivity.

The PPI matrices of the Go or Stop conditions from the “generalized” PPI approach were very different from the BSC matrices of respective conditions. This is because the PPI effects for one condition reflect connectivity differences between the very condition and the rest of the experiment period. This cannot be achieved by using the BSC method, because the implicit baseline conditions cannot be easily modeled as events. Compared with the implicit baseline, the Go and Stop conditions showed decreased connectivity between visual areas, and increased connectivity between visual areas and sensorimotor areas among other brain regions. The reduced connectivity within the visual areas during task execution compared with baseline is consistent with our previous studies using a set of different tasks (Di et al., 2017) as well as in a simple checkerboard task (Di & Biswal, 2017). However, in contrast to the reduced functional connectivity between visual and sensorimotor regions in the checkerboard task (Di & Biswal, 2017), the current results showed increased functional connectivity between the visual and sensorimotor regions. It is not surprising because the stop signal task requires the subjects to response to visual stimuli, therefore yielding increased functional coupling between visual and sensorimotor regions.

When directly comparing the differences between the Stop and Go conditions, both PPI and BSC methods showed similar results, which are consistent with our explanations in the introduction. In addition to Spearman’s correlation used in our previous paper (Di & Biswal, 2019), we further compared BSC differences using Pearson’s correlation, covariance, and beta series-by-task interaction in the current analysis. Pearson’s correlation did yield more statistical significant effects than Spearman’s correlation, however, all the four measures showed less number of significant effects than the PPI analysis. Therefore, in terms of statistical sensitivity, there is no support for the claim that BSC is more suitable for event-related designed data (Cisler et al., 2014).

In this paper, we have explained the deconvolution process for the BOLD signals, which can help to understand the relations between PPI and BSC. The success of deconvolution is critical for PPI and BSC analyses on event-related designed data. The different ways of dealing with deconvolutions or single trial estimations may partly contribute to the different results between PPI and BSC analyses. Further studies may seek to improve the deconvolution process to improve both PPI and BSC. For example, more sophisticated filters could be used for deconvolution, e.g. cubature Kalman filtering (Havlicek, Friston, Jan, Brazdil, & Calhoun, 2011). In addition, applying subject-specific or region-specific HRF (Pedregosa, Eickenberg, Ciuciu, Thirion, & Gramfort, 2015) may also be helpful given the large variability of HRFs (Handwerker et al., 2004). Lastly, optimizations of event-related design may also improve the deconvolution process and single trial estimates, thus providing better connectivity estimates for both methods (Abdulrahman & Henson, 2016; Mumford et al., 2014; Visser et al., 2016).

## 5. Conclusion

In the current paper, we explained how PPI and BSC measure task modulated connectivity. PPI is a model based approach to examine differences in effective connectivity in difference task conditions. In contrast, a BSC measures correlations of observed trial-by-trial activations in a certain condition. However, when comparing the differences between task conditions, BSC should in principle yield similar results as shown by PPI.

## Author contributions

X.D. conceived the idea, performed the data analysis, and wrote the draft. All authors discussed the results, and contributed to the final manuscript.

## Compliance with Ethical Standards

This study involves re-analysis of an open-access fMRI dataset. We did not use any personal identifiable information in the current analysis.

