## Supplementary materials for "Understanding psychophysiological interaction and its relations to beta series correlation"

**Supplementary materials for “Psychophysiological interaction and beta series correlation for task modulated connectivity: modeling considerations and their relationships”**

Xin Di ^1, 2^, Zhiguo Zhang ^3, 4^, Bharat B Biswal ^1, 2 *^

1, School of Life Sciences and Technology, University of Electronic Science and Technology of China, Chengdu, China

2, Department of Biomedical Engineering, New Jersey Institute of Technology, Newark, NJ, 07029, USA

3, School of Biomedical Engineering, Health Science Center, Shenzhen University, Shenzhen, China

4, Guangdong Provincial Key Laboratory of Biomedical Measurements and Ultrasound Imaging, Shenzhen, China

This document includes supporting information to the manuscript titled “Psychophysiological interaction and beta series correlation for task modulated connectivity: modeling considerations and their relationships”.

Contents:

Supplementary 1: Centering will not alter the interpretation of a variable

Supplementary 2: Simulation results

Supplementary 3: The effects of event duration setting on the PPI results

Supplementary 4: The asymmetry of PPI matrix

**Supplementary 1: Centering will not alter the interpretation of a variable**

In this section, we show that if the constant term is included in the model, centering of other variables (removing the mean) will not affect the effect estimates of these variables.

Given an observed variable *y*, we could fit a linear model using a psychological variable *x_Psych_*. A set of *β* estimates of *β_0_* and *β_1_* could be obtained from the model A1.

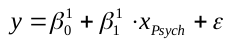
 A1

Now we define another psychological variable *x^*^_Psych_*, where it contains an additional constant value *c*.

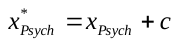
 A2

We could fit *y* with another linear model with the new psychological variable *x^*^_Psych_*, and obtain a new set of *β* estimates.

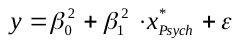
 A3

Substitute *x^*^_Psych_*, and we can have the following:

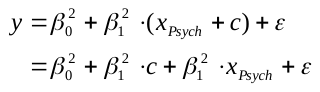
 A4

Compare A1 and A4, we can see the following relationship:

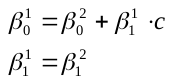
 A5

That is, because the constant term is already in the model, adding a constant component to the *x_Psych_* variable will not affect the estimated effect of *x_Psych_*.

**Supplementary 2: Simulation results**

We have explained in the main text that for the two models in Figure 1A and 1B (also Figure S1A and S1D), although the first regressor in model A (A_1_) and the second regressor in model D (D_2_) are identical, because of the differences in the other regressors in the models, the meanings of the regressors A_1_ and D_2_ are different. A_1_ from the first model represents the condition specific effect of the first condition, while D_2_ from the second model represents the differences between the two conditions. We simulated 1,000 times using Gaussian random variables with the mean of the first condition as 1 and the mean of the second condition as a random variable. We used the two models to fit the data to obtain the effects of the regressors A_1_ and D_2_ from the two models. We plotted the effect estimates of the regressors A_1_ and D_2_ against the averaged effects of the first and second conditions, respectively in Figure S1. It shows clearly that the effect estimates of A_1_ were only correlated with the mean of the first condition (Figure S1B), but not correlated the second condition (Figure S1C). In contrast, the effect estimates of D_2_ were not correlated with the mean of the first condition (Figure S1E), but reversely correlated the mean of the second condition (Figure S1F). This suggests that D_2_ reflects the differences between the two conditions rather than the specific effect of the first condition. The code for this simulation is available at <https://github.com/dixy0/PPI_Intro/blob/master/Condition_modeling.m> .

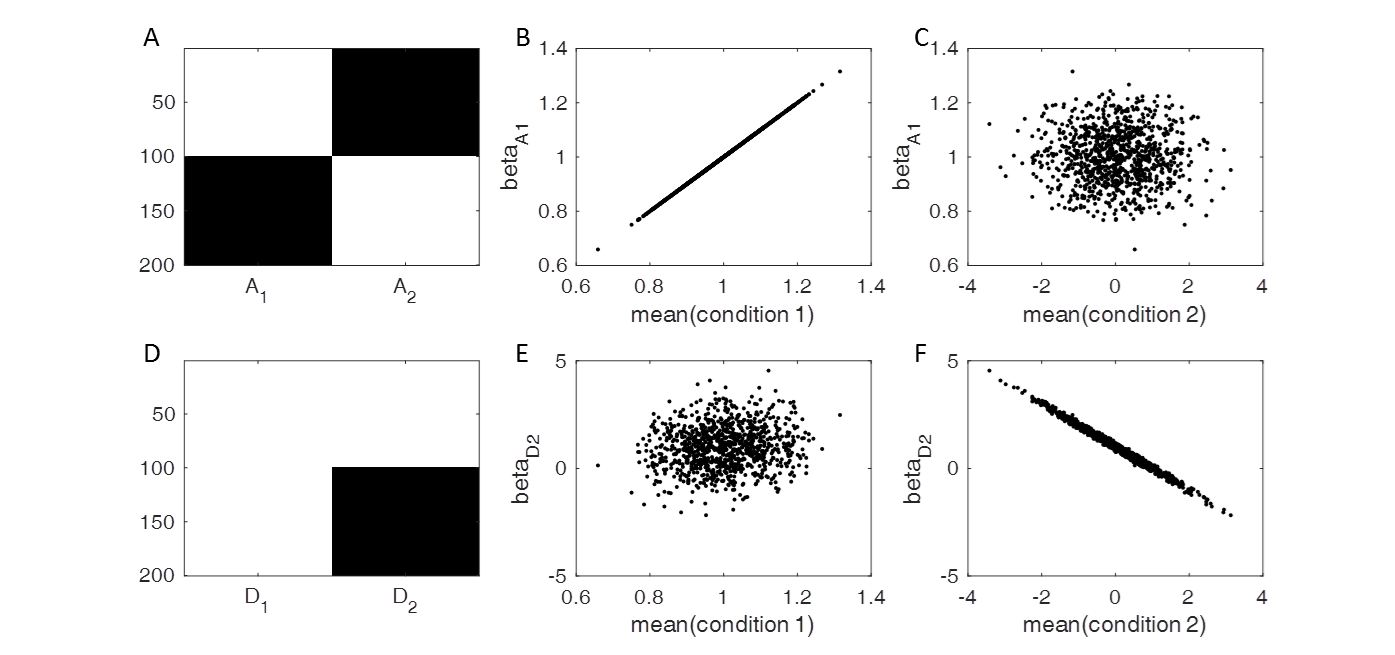

**Figure S1**

Similarly in the interaction models, although the interaction term PPI_A1_ in the model depicted in Figure S2A is the same as the PPI term PPI_D2_ in the model depicted in Figure S2D (Figure 1C and 1D in the main text), because the differences of the other interaction terms, the meaning of PPI_A1_ and PPI_D2_ from the two models are different. PPI_A1_ from the first model represents the condition specific relationships with seed in the first condition, while PPI_A2_ from the second model represents the different relationships with the seed between the two conditions. We simulated 1.000 times using Gaussian random variables with a fixed relationship between the seed and test variables in the first condition and a variable relationship in the second condition. We used the two models to fit the data to obtain the effects of PPI_A1_ and PPI_D2_. We plotted the effect estimates of PPI_A1_ and PPI_D2_ against the correlations between the seed and tested variables in the first and second conditions, respectively. It shows clearly that the effect estimates of PPI_A1_ are only correlated with the correlations in the first condition (Figure S2B), but not with the correlations in the second condition (Figure S2C). In contrast, the effect estimates of PPI_D2_ are not correlated with the correlations in the first condition (Figure S2E), but reversely correlated the correlation in the second condition (Figure S2F). This suggests that PPI_D2_ reflects the differential relationships between the two conditions rather than the specific correlation in the first condition. The code is available at <https://github.com/dixy0/PPI_Intro/blob/master/Interaction_modeling.m> .

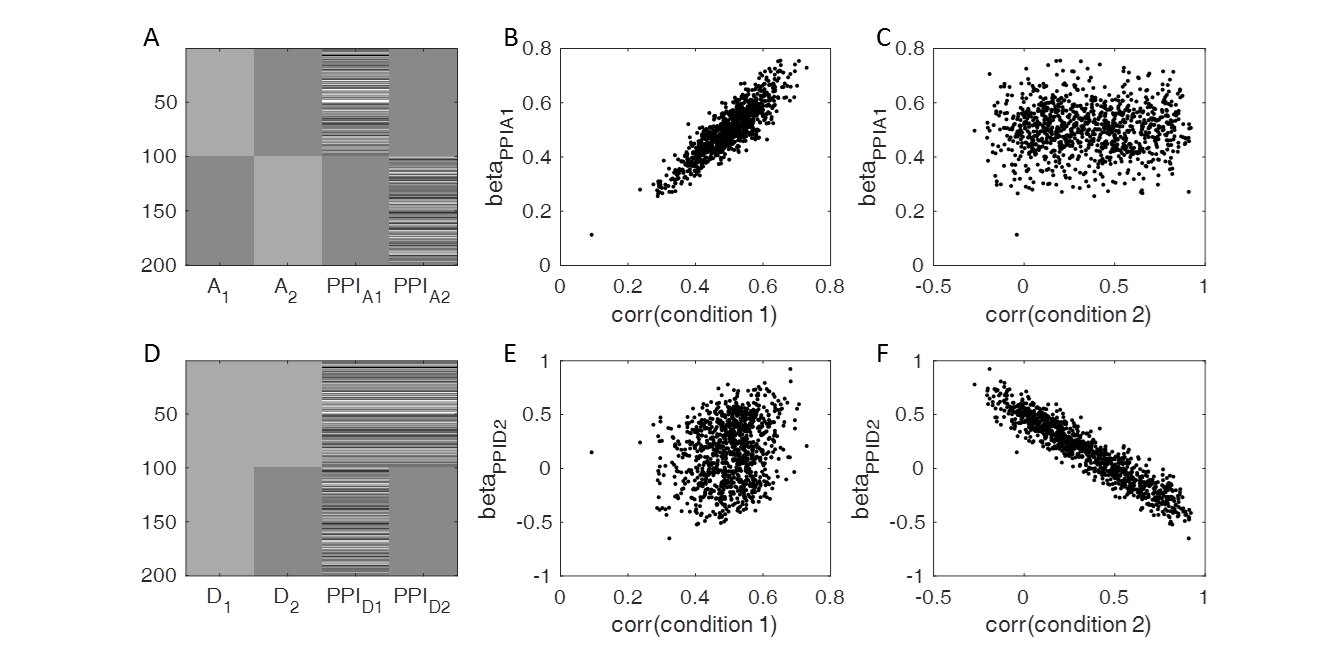

**Figure S2**

**Supplementary 3: The effects of event duration setting on the PPI results**

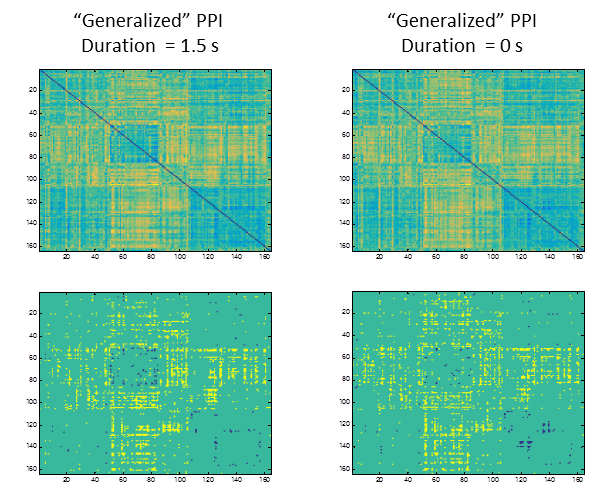

**Figure S3** The effect of event duration setting on the PPI results. When setting GLM models for defining PPI terms, the duration of events for a event-related design can be modeled as 0 to represent a impulse of evoked neuronal responses or the actually duration of the trials. This duration setting may alter the shape of the PPI terms, thus altering the PPI results. Here, however, we show that if we used the duration of 0 for calculating the PPI terms, the PPI results for the Stop vs. Go conditions in the stop signal task were very similar to those using the actual trial duration of 1.5 s, which was used in the main analysis.

**Supplementary 4: The asymmetry of PPI matrix**

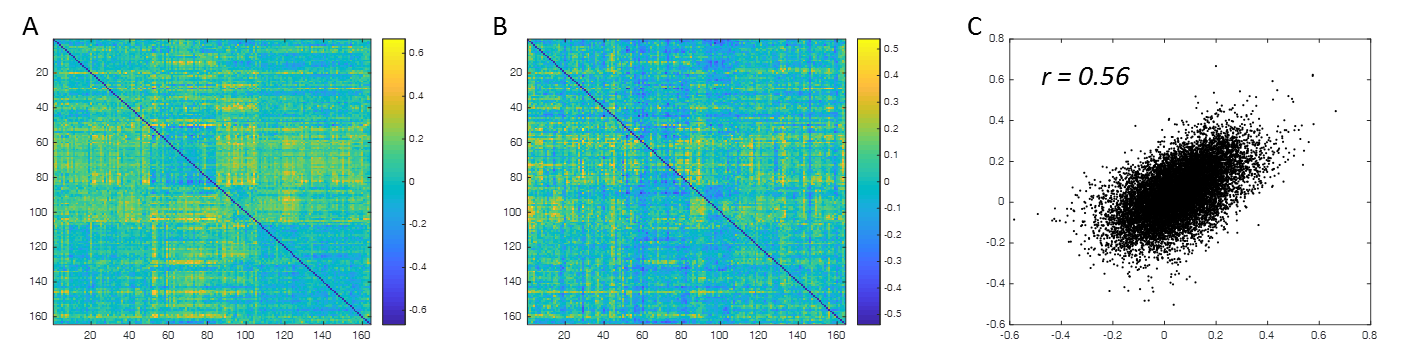

**Figure S4** The asymmetry of the PPI matrix corresponding to the contrast between the Stop and Go conditions. A) Mean PPI matrix of the contrast between Stop and Go condition without symmetrization. B) Differential matrix between the raw and transposed PPI matrices. C) Scatter plot of the corresponding PPI effects of the lower and upper diagonals in matrix A. The Pearson’s correlation between the lower and upper diagonals was *0.56*.
